# Sign-consistency based variable importance for machine learning in brain imaging

**DOI:** 10.1101/124453

**Authors:** Vanessa Gómez-Verdejo, Emilio Parrado-Hernández, Jussi Tohka, Alzheimer’s Disease Neuroimaging Initiative

**Author notes:** Data used in preparation of this article were obtained from the Alzheimers Disease Neuroimaging Initiative (ADNI) database (adni.loni.usc.edu). As such, the investigators within the ADNI contributed to the design and implementation of ADNI and/or provided data but did not participate in analysis or writing of this report. A complete listing of ADNI investigators can be found at adni.loni.usc.edu/wpcontent/uploads/how_to_apply/ADNI_Acknowledgement_List.pdf.

## Abstract

An important problem that hinders the use of supervised classification algorithms for brain imaging is that the number of variables per single subject far exceeds the number of training subjects available. Deriving multivariate measures of variable importance becomes a challenge in such scenarios. This paper proposes a new measure of variable importance termed sign-consistency bagging (SCB). The SCB captures variable importance by analyzing the sign consistency of the corresponding weights in an ensemble of linear support vector machine (SVM) classifiers. Further, the SCB variable importances are enhanced by means of transductive conformal analysis. This extra step is important when the data can be assumed to be heterogeneous. Finally, the proposal of these SCB variable importance measures is completed with the derivation of a parametric hypothesis test of variable importance. The new importance measures were compared with a t-test based univariate and an SVM-based multivariate variable importances using anatomical and functional magnetic resonance imaging data. The obtained results demonstrated that the new SCB based importance measures were superior to the compared methods in terms of reproducibility and classification accuracy.

## 1 Introduction

Machine Learning (ML) is a powerful tool to characterize disease related alterations in brain structure and function. Given a training set of brain images and the associated class information, here a diagnosis of the subject, supervised ML algorithms learn a voxel-wise model that captures the class information from the brain images. This has direct applications to the design of imaging biomarkers, and the inferred models can additionally be considered as multivariate, discriminative representations of the effect of the disease to brain images. This representation is fundamentally different from conventional brain maps that are constructed based on a voxel-by-voxel comparison of two groups of subjects (patients and controls) and the patterns of important voxels in these two types of analyses provide complementary information (Kerr et al 2014; Haufe et al 2014; Tohka et al 2016).

A key problem in using voxel-based supervised classification algorithms for brain imaging applications is that the dimensionality of data (the number of voxels in the images of a single subject, i.e., the number of variables^1^) far exceeds the number of training subjects available. This has led to a number of works studying variable selection within brain imaging; see Mwangi et al (2014) for a review. However, in addition to selecting a set of relevant variables, it is interesting to rank and study their importance to the classification. This problem, termed variable importance determination, has received significantly less attention and is the topic of this paper.

The simplest approach to assess the importance of a variable is to measure its correlation with the class labels, for example, via a t-test. This is exactly what massively univariate analysis does. However, it considers variables independently of others and, therefore, may miss interactions between variables. Indeed, a variable can be meaningful for the classification despite not presenting any linear relationship with the class label (Haufe et al 2014). Further, there is evidence that this importance measure does not perform well for variable selection in discrimination tasks (Chu et al 2012; Tohka et al 2016) and, therefore, multivariate importance measures might be more appropriate.

The use of ML opens a door for more sophisticated methods to assess the importance of variables, able to capture some of the interactions among variables. A straightforward approach is to base each variable importance on the difference between the performances observed in an ML method when the variable is present and when it is absent from the training instances. A significant decrease in performance after removing a variable indicates its relevance. While this methodology is straight-forward, it is not suitable for brain imaging applications as it would fail to recognize the relevance of important but mutually redundant variables. A few ML methods favor a richer and more detailed analysis as they enable to assess the role that each individual variable plays in the composition of the architecture of the model. Two clear examples of this class are linear models and Random Forests^2^ (RFs).

If the variables have been properly standardized, the weights of a linear classifier can be considered as measures of variable importance (Caragea et al 2001) (see, e.g, Cohen et al (2010); Khundrakpam et al (2015) for neuroimaging examples). However, when the variables are redundant as often in neuroimaging, the weights may change erratically in response to small changes in the model or the data. This problem is known as multicollinearity in statistics. Linear regressors can be endowed with Lasso and Elastic Net regularizations (Friedman et al 2008; Zou and Hastie 2005), in order to deal with problems with a high number of input variables. These regularizations force sparsity and remove variables of reduced relevance from the linear model, enhancing the contribution of the remaining variables. More elaborated methods take a further step in the exploitation of the relationship between the weight of each variable in a linear classifier/regressors and its relevance (Guyon et al 2002). The starplots method of Bi et al (2003) learns an ensemble of linear Support Vector Regressors (SVR) endowed with a Lasso type regularization in the primal space. The regularization filters out the non-relevant variables from each regressor, while the starplots determine the relevance of the non-filtered variables by looking for smooth and consistent patterns in the weights they achieve across all the regressors in the ensemble. These linear methods with Lasso regularization present two significant drawbacks in very high dimensional scenarios. First, the computational burden of the resulting optimization in the primal space is high. Second, they enforce an aggressive sparsity, usually reducing the number of non-filtered variables to a final quantity comparable to the number of training instances. This last feature is specially harmful in neuroimaging problems, where the typical situation is that a disease affects a set of focal brain areas. This reflects in groups of clustered variables being important with a strong correlation. A Lasso regularization would filter out most of the important (but correlated) variables what complicates the interpretation of the discrimination pattern. To combat this problem, for example, Grosenick et al (2013) and Michel et al (2011) have introduced brain imaging specific regularizers which take into the account the spatial structure of the data. The application of these methods is complicated by a challenging parameter selection (Tohka et al 2016) and deriving a variable-wise importance measure is complicated by the joint regularization of weights of the different variables.

Also RFs (Breiman 2001) facilitate the assessment of variable importance. RFs are ensembles of decision trees where each tree is trained with a subset of the available training subjects, and with a subset of the available variables. RFs offer two main avenues for assessing the variable importance: Gini importance and permutation importance based on the analysis of out-of-bag samples (Archer and Kimes 2008). Both measures have found applications in brain imaging: Langs et al (2011) studied voxel selection based on Gini importance, Moradi et al (2015) ranked the different types of variables (imaging, psychological test scores) for MCI-to-AD conversion prediction based on the out-of-bag variable importance and Greenstein et al (2012) ranked the importance of cortical ROI volumes to schizophrenia classification. However, these applications have considered at most tens of variables while our focus is on a voxel-wise analyses of whole brain scans, where we have tens of thousands variables. Indeed, the usability of RFs to capture variable importance in a multivariate fashion is questionable in high dimensional scenarios. This is partially due to the use of decision trees as a base classifiers. Each decision tree in an RF comes out of a training set that includes a sample of the observations and of the variables. In a data set in which the number of variables is far larger than the number of observations each tree definition will rely on a very reduced set of variables (notice that a set of 100 samples is split completely by a tree with 100 nodes, each implementing a threshold test in one of the variables). This means that in order to get a chance to assess the importance of all the variables (namely, that each variable shows a large enough number of times across the trees in the ensemble), the forest must contain an extraordinarily large number of trees, and this makes the method computationally less attractive than the use of an ensemble of linear classifiers. In an ensemble of linear classifiers every variable can get a weight in every classifier, while in a random forest typically only a reduced fraction of variables will be used as splits in every tree. In addition, the weight of a variable in a linear classifier results from a global optimization process that takes into account the joint contributions of all variables simultaneously, while the usage of a variable in a split within a decision tree is strongly dependent of the particular subset of splits that lead to the branch in which this split is used (Strobl et al 2008). Note that this criticism pertains only to the variable importance scores from RFs, and not to the accuracy of predictions derived from RFs.

To overcome the limitations of the regularized linear models and RFs, we introduce and study a new variable importance measure based on sign consistency of the weights in an ensemble of linear Support Vector Machines (SVMs). Briefly, we train an ensemble of SVMs using only a part of the subjects available for each SVM in the ensemble. The main idea is to define the importance of a variable using its sign consistency, i.e., the fraction of members of the ensemble in which its weight is positive (or negative). We thereafter prune the variable importances using the ideas from transductive conformal analysis. We derive parametric hypothesis test of the variable importance measures and show that the new importance measures are an improvement over p-value estimation for SVM weights of Gaonkar and Davatzikos (2013); Gaonkar et al (2015).

The results presented in this paper build on our previous work on variable selection (Parrado-Hernández et al 2014) and thoroughly extend a preliminary conference paper focused on assessing variable relevance (Gomez-Verdejo et al 2016). In particular, the main novel contributions of this paper over (Parrado-Hernández et al 2014) and (Gomez-Verdejo et al 2016) are:

- We explain the variable importance measure and its properties in more detail than before and provide algorithms for its computation.
- We derive a parametric hypothesis test for variable importance. The variable selection is carried out with a hypothesis test that replaces the cross validation of the previous works. The hypothesis test improves the stability and reduces the processing time by an order of magnitude.
- We perform new large-scale experiments to assess the accuracy and stability of our novel importance measure in comparison to related approaches (Gaonkar and Davatzikos 2013; Gaonkar et al 2015) and show that this new measure is an improvement over them.

## 2 Methods

### 2.1 Variable importance with ensembles of linear SVMs

We start by introducing the basic variable importance computation, termed Sign Consistency Bagging (SCB). Thereafter, in the next subsection, we enhance the basic algorithm by introducing a conformal variant (SCBconf) of the basic algorithm that outperforms the basic algorithm for certain problems.

Consider a binary classification task to differentiate between two groups of subjects. We refer to these two groups as patients and controls. The training data is formed by *N* pairs (**x**^*i*^*, y*^*i*^), *i* = 1*,…, N* where 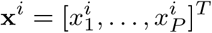 is a vector with the variables corresponding to the brain scan of the *i*-th subject and *y*_*i*_ = 1 if the *i*-th subject is a patient and *y*_*i*_ = *-*1 otherwise. We assume 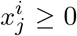. This is often a natural requirement and, in any case, it can be fulfilled by adding a suitable constant to all 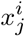.

We use linear SVM classifiers as base learners. The predicted label *ŷ* for a test sample **x** is given by

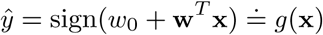

and the classifier is defined in terms of its parameters *w*_0_ (bias) and **w** = [*w*_1_*,…, w*_*P*_]^*T*^ (weight vector).

We train *S* linear SVMs, each with a different subset of *γN* training instances selected at random without replacement; *γ* ∈ [0, 1] is the subsampling rate, usually selected as *γ* = 0.5. The *s*-th SVM is therefore described by the bias term *w*^*s*^ and the weight vector 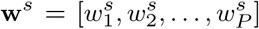. Once the learning of the ensemble is finished, each variable in the input data *x*_*j*_ is related with a set of *S* weights 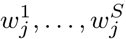. Since 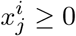 for all *i, j*, there is a straightforward qualitative interpretation of the sign consistency across all the 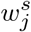 in the set. If 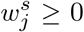 for all *s*, we have that the *j*-th variable pushes the classifier towards the positive class while a result of 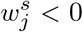 for all *s* means that *j*-th variable pushes the classifier towards negative class. Since the linear SVM uses a *L*_2_ norm regularization that does not enforce sparsity in the primal space it is very unlikely that 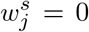, for all *j, s*. It is rare that all 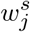 would share the same sign and we introduce an importance scoring *I*_*j,*_ *j* = 1*,…, P,* quantifying the sign consistency:

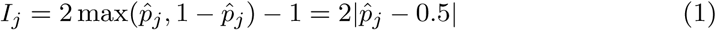

with

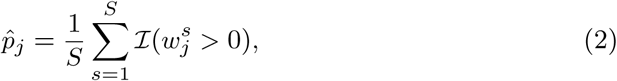

where *ℐ*(*x*) is the indicator function, *ℐ*(*x*) = 1 if *x* is true and *ℐ*(*x*) = 0 otherwise. If all weights corresponding to a same variable 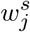*, s* = 1*,…, S* receive the same sign, variable *j* would get an importance scoring *I*_*j*_ = 1, meaning it is very relevant for the classification. As the proportion of 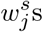 with opposite sign get balanced, the interpretation of the variable *j* for the classification gets blurred, since in some members of the ensemble a high value of *x*_*j*_ would indicate member of one of the classes while in some other members it would indicate membership of the other class. In this case, as the interpretation varies depending on the sub-sample, the variable is not useful (or is even harmful) for the generalization performance and should have a low importance score.

### 2.2 Transductive refinement of variable importance

In this sub-section, we introduce a variant of SCB variable importance, SCBconf. This algorithm builds on the results of SCB and enhances the identification of the relevant variables by borrowing ideas from transductive learning and conformal analysis.

Transduction refers to learning scenarios in which one has access to the observations, but not the labels, of the test set^3^. Conformal analysis goes a step further and proposes to learn a set of models, one per each training set that arises from adding the test instance with each possible label (in binary classification one would learn two models). Then, the test instance is classified with the label that led to the model that better conformed to the data.

Conformal analysis is used to enhance variable importance scores as follows: Let 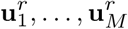 be a subset of *M* testing data selected randomly in the *r*-th conformal iteration with *r* = 1*,…, R*. Now, *M* independent labellings 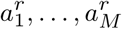 are generated at random. Label 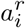 is the one generated for sample 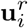 in the *r*-th iteration. The correct labels of these test samples are never used along this procedure because they are not accessible. For each of these iterations, we compute the importance measures *I*_*j*_ (*r*), *j* = 1*,…, P,* based on the training data **x**_1_*,…,* **x**_*N,*_ the test samples **u**_1_*,…,* **u**_*M*_ and the labels 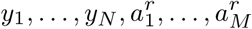. After running R iterations, we set

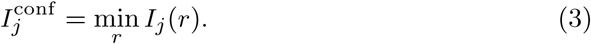

For a variable to be important, 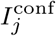 requires that it is important in all of the *R* labellings. The underlying intuition is that the importance of variables that yield a high *I*_*j*_ (*r*) in a few the subsets, but not in all of them, strongly depends on particular labellings. Therefore these variables should not be selected as their importances are not aligned with the labeling that leads to the disease discrimination, but labellings that stress other partitions not relevant for the characterization of the disease.

### 2.3 Hypothesis test for selecting important variables

The previous subsections have introduced two scorings, *I*_*j*_ and 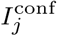, able to assess the relevance of the variables. This subsection presents a hypothesis test to fix qualitative thresholds so that variables with scorings above the threshold can be considered as relevant for the classification and variables with scorings below the threshold can be safely discarded since their importance is reduced. We first present the test for importances *I*_*j*_ and thereafter generalize this test for 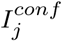 We adopt a probabilistic framework in which the sign of the weight of variable *j* in the SVM of bagging iteration *s*, 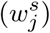, follows a Bernoulli distribution with the unknown parameter *p*_*j*_ *∈* (0, 1); this indicates that *w*_*j*_ *>* 0 with probability *p*_*j.*_ In this framework, an irrelevant variable *j* is expected to yield positive and negative values in *w*_*j*_ with the same probability, thus one would declare variable *j* as irrelevant if *p*_*j*_ = 0.5 with high probability. We formulate this scenario via the following hypothesis test:

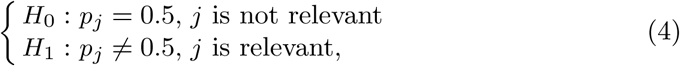

which we use to detect relevant variables by rejecting the null hypothesis. We solve the test (4) with a statistic *z*_*j*_:

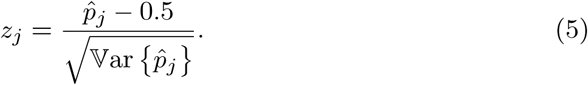

where estimate 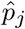 of *p*_*j*_ is computed as the sample mean of the observed signs of 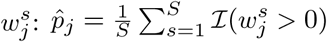. The parameter 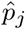 can be considered to follow a binomial distribution rescaled by the factor *S* and, thus, its variance can be estimated as 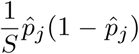 from the observations. As the observations come from a bagging process, they are correlated and independence cannot be assumed. Therefore, we resource to the following estimator of 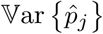(Nadeau and Bengio 2003):

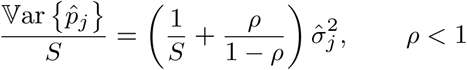

where *ρ* represents the correlation among samples and 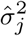 denotes to the estimator variance if independence could be assumed. Moreover, according to Nadeau and Bengio (2003), since the bagging corresponds to a scenario in which, at each iteration, *n*_1_ samples are used for training the SVM and *n*_2_ = *N - n*_1_ are left out, *ρ* can be estimated as *n*_1_*/*(*n*_1_ + *n*_2_). Since the proposed bagging scheme uses *n*_1_ = *γN* training samples in each iteration, we can approximate *ρ* with *γ* and, noticing also that *S* ≫ 1, we get that

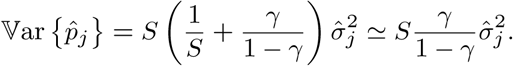

With these approximations, the statistic *z*_*j*_ of (5) becomes

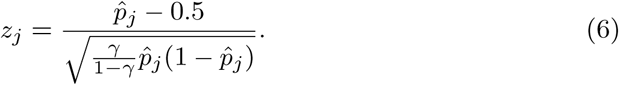

The statistic *z*_*j*_ of (6) follows a t-student distribution with *S -* 1 degrees of freedom (Nadeau and Bengio 2003). When *S* is large enough, as in our case, one scan safely approximate the distribution by the standard Gaussian distribution with zero mean and unit variance.

We now generalize the hypothesis test to the importances 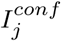 of Subsection 2.2. The selection of 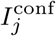 as the minimum of the *R* scorings *I*_*j*_ (*r*) is equivalent to select as 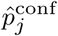 the 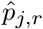 that lies closest to 0.5. The 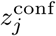 can be then computed using Eq. (6) and substituting 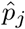 by 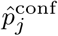. An equivalent definition would be to select *z*_*j*_ with the smallest absolute value among the *R* candidates.

#### Algorithm 1 Sign Consistency Bagging

**Input:** *X*: *N* × *P* matrix with training brain scans (each row is a subject, each column a variable); **y**: vector with the labels corresponding to the rows of *X*

**Output:** *I*: *P* × 1 vector with voxel relevances; **z**: *P* × 1 vector with significance statistic

1: 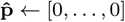 vector with *P* zeros

2: **for** *s* = 1 to *S* **do**

3: *X_s_,* **y***_s_* ← randomly sample *γN* training samples

4: **w**^*s*^ ← LinearSVM(*Xs,* **y***s*)

5: **for** *j* = 1 to *P* **do**

6: **if 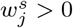 then**

7: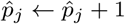

8: *I* = []

9: **z** = []

10: **for** *j* = 1 to *P* **do**

11: 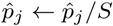

12: *I*_*j*_ ← 2 max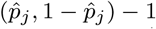

13: Compute score *z*_*j*_ using (6)

#### Algorithm 2 Sign Consistency Bagging with transductive refinement

**Input:** *X*: *N*× *P* matrix with training brain scans (each row is a subject, each column a variable); **y**: vector with the labels corresponding to the rows of *X*; *X_t_*: *Nt* × *P* matrix with testing brain scans

**Output: I**^conf^: *P* × 1 vector with variable relevances; **z**: *P* × 1 vector with significance statistic

1: *I*← [] empty *P* × *R* matrix

2: **for** *r* = 1 to *R* **do**

3: *U*^*r*^ ← randomly sample *M* testing observations from matrix *Xt*

4: *A*^*r*^ ← randomly generate a label 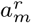 per each 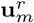*, m* = 1*,…, M*

5: 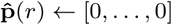vector with *P* zeros

6: **for** *s* = 1 to *S* **do**

7: *X_s_,* **y***_s_* ← randomly sample *γN* training data

8: 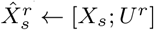

9: 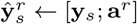

10: **W**^*s*^ ← LinearSVM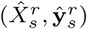

11: **for** *j* = 1 to *P* **do**

12: **if 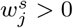 then**

13: 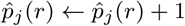

14: **for** *j* = 1 to *P* **do**

15: 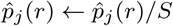

16: *I*_*j*_(*r*) ← 2 max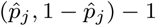

17: **I**^conf^ ← [] empty vector with *P* elements

18: **for** *j* = 1 to *P* **do**

19: 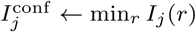

20: *b* ← arg min_*r*_ *I*_*j*_(*r*)

21: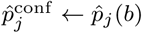

22: Compute score *z*_*j*_ using (6) and 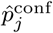

### 2.4 Implementation

Algorithms 1 and 2 sketch the implementation of the method to assess variable importance and its version with transductive refinement. In both cases, the ensemble contains a total of *S* = 10.000 SVMs and each SVM is trained with half of the available training data (*γ* = 0.5). The SVM regularization parameter *C* was fixed to 100 as it was observed to be large enough to solve properly these linearly separable problems. A supplement further demonstrates the use of the SCB method with a small toy example.

If any of the training sets presents unbalanced class proportions, the subsampling process at each bagging iteration is used to correct for it. If the transductive refinement is applied, the number of conformal iterations is set to *R* = 20. For each of these iterations, the number of selected test data, *M,* has been fixed in such a way that no more than two test data samples is used per each 100 training samples. The hypothesis test described in Subsection 2.3 to identify the subset of important variables is applied with a significance level of *α* = 0.05.

Finally, the overall goodness of the proposed variable importance measure is evaluated by checking the discriminative capabilities of a linear SVM trained using only the important variables. This SVM is also trained with *C* = 100, since in most cases there are still more variables than samples. However, unlike in the bagging iterations, in this final classifier the class imbalance is solved by using a reweighting the regularization parameter of the samples of the minority class in the training of the SVM. This way the contribution of the samples of both minority and majority class to the SVM loss function is equalized. This is a standard procedure within SVMs, contained in most SVM implementations (Chang and Lin 2011).

The software implementation of all the methods has been developed in Python^4^. The SVM training relies on the Scikit-learn package (Pedregosa et al 2011) which is based on the LIBSVM (Chang and Lin 2011).

## 3 Materials

### 3.1 Simulated data

We generated 10 simulated data sets to evaluate the methods against the known ground-truth and to demonstrate their characteristics with a relatively simple classification task. The datasets contained simulated images of 100 controls and 100 patients and the images had 29852 voxels, similarly to ADNI magnetic resonance imaging (MRI) data in the next subsection.

The simulations were based on the AAL atlas (Tzourio-Mazoyer et al 2002), downsampled to 4*mm*^3^ voxel-size. We selected six regions as important modeling dementia related changes in structural MRI. The voxels of these regions are given in sets *Q*_1_*,…, Q*_6_ which are left and right Hippocampus (*Q*_1_*, Q*_2_), Thalamus (*Q*_3_*, Q*_4_), and Superior Frontal Gyrus (*Q*_5_*, Q*_6_). We generated the brain scan corresponding to the each subject in a way that each relevant region (*Q*_1_*,…, Q*_6_) presents correlated voxels (within a class), to make the task of finding them difficult for multivariate variable selection/importance methods. For every subject *i*, *i* = 1*,…, N,* we draw a subject bias *b*_*i*_ from a Gaussian distribution with zero mean and variance 0.01. Define *Q* as the union of the six relevant regions *Q*_*k.*_ If *i* models a patient, its voxel values are generated with the rule:

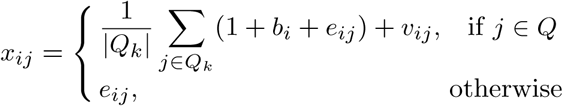

If *i* models a control subject, its voxel values are generated with:

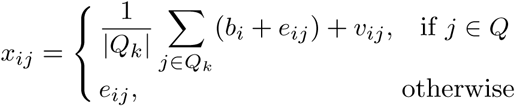

In all cases *e*_*ij*_ and *v*_*ij*_ were drawn from zero-mean Gaussian distributions with variances 1 and 0.01, respectively. Thereafter, to the relevant voxels, we added white noise with the variance 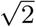 projected to the Bayes-optimal decision hyperplane. This operation maintains the Bayes error rate, but it makes the task of finding important voxels more difficult. Finally, we filtered the images with an isotropic 4-mm FWHM Gaussian kernel to model the smoothness in brain images. The Bayes error for this data was 2.2 %. To evaluate the classification accuracy of the methods, we simulated a large test set with the same parameters as the training set.

### 3.2 ADNI data

A part of the data used in the preparation of this article were obtained from the Alzheimer’s Disease Neuroimaging Initiative (ADNI) database (adni.loni.usc.edu). For up-to-date information, see www.adni-info.org.

We studied the classification between MCI (mild cognitive impairment) and healthy subjects (NCs) using structural MRIs from ADNI. This problem is more challenging than NC vs. Alzheimer’s Disease (AD) classification (Tohka et al 2016) and therefore offers better insight into the capabilities for different variable importance methods. We did not consider stable vs. progressive MCI classification as the number of MCI subjects is not large enough for the reproducibility analysis performed with this data (see Tohka et al (2016) for a more detailed discussion). We used MRIs from 404 MCI subjects and 231 NCs (T1-weighted MP-RAGE sequence at 1.5 Tesla, typically 256 x 256 x 170 voxels with the voxel size of 1 mm x 1 mm x 1.2 mm). The MRIs were preprocessed into gray matter tissue images in the stereotactic space, as described in Gaser et al (2013); Moradi et al (2015), and thereafter they were smoothed with the 8-mm FWHM Gaussian kernel, resampled to 4 mm spatial resolution and masked into 29852 voxels. We age-corrected the data by regressing out the age of the subject on a voxel-by-voxel basis (Moradi et al 2015). This has been observed to improve the classification accuracy in dementia related tasks (Tohka et al 2016; Dukart et al 2011) due to overlapping effects of normal aging and dementia on the brain.

With these data, we studied the reproducibility of the variable importance using split-half resampling (aka 2-fold cross-validation) akin to the analysis in (Tohka et al 2016). We sampled without replacement 100 subjects from each of the two classes, NC and MCI, so that *N* = 200. This procedure was repeated *L* = 100 times. We denote the two subject samples (split halves; train and test) by *A*_*l*_ and *B*_*l*_ for the iteration *l* = 1*,…, L*. The sampling was without replacement so that the split-half sets *A*_*l*_ and *B*_*l*_ were always disjoint and therefore can be considered as independent train and test sets. The algorithms were trained on the split *A*_*l*_ and tested on the split *B*_*l*_ and, vice versa, trained on *B*_*l*_ and tested on *A*_*l*_. All the training operations, including the estimation of regression coefficients for age removal, were done in the training half. The test half was used only for the evaluation of the algorithms. We used 2-fold CV instead of more typical 5 or 10-fold CV as we needed to ensure that the training sets between different folds are independent in order not to overestimate the reproducibility of variable importance scores.

### 3.3 COBRE data

To demonstrate the applicability of the method for the resting state fMRI analysis, we used the pre-processed version^5^ of the COBRE sample (Bellec et al 2015). The dataset, which is a derivative of the COBRE sample found in International Neuroimaging Data-sharing Initiative (INDI)^6^ includes preprocessed resting-state fMRI for 72 patients diagnosed with schizophrenia (58 males, age range = 18-65 yrs) and 74 healthy controls (51 males, age range = 18-65 yrs). The fMRI dataset features 150 EPI blood-oxygenation level dependent (BOLD) volumes (TR = 2 s, TE = 29 ms, FA = 75 degrees, 32 slices, voxel size = 3×3×4 *mm*^3^, matrix size = 64×64) for each subject.

We processed the data to display voxel-wise estimates of the long range functional connectivity (Guo et al 2015). It is well documented that a disruption of intrinsic functional connectivity is common in schizophrenia patients, and this disruption depends on connection distance (Wang et al 2014; Guo et al 2015). First, the fMRIs were preprocessed using the NeuroImaging Analysis Kit (NIAK^7^) version 0.12.14 as described at ^4^. The preprocessing included slice timing correction and motion correction using a rigid-body transform. Thereafter, the median volume of fMRI of each subject was coregistered with the T1-weighted scan of the subject using the Minctracc tool (Collins and Evans 1997). The T1-weighted scan was itself non-linearly transformed to the Montreal Neurological Institute (MNI) template (symmetric ICBM152 template with 40 iterations of non-linear coregistration (Fonov et al 2011)). The rigid-body transform, fMRI-to-T1 transform and T1-to-stereotaxic transform were all combined, and the functional volumes were resampled in the MNI space at a 3 mm isotropic resolution. The “scrubbing” method (Power et al 2012) was used to remove the volumes with excessive motion (frame displacement greater than 0.5 mm). A minimum number of 60 unscrubbed volumes per run, corresponding to 180 s of acquisition, was required for further analysis. For this reason, 16 controls and 29 schizophrenia patients were rejected from the subsequent analyses, yielding 43 patients and 58 healthy controls to be used in the experiment. The following nuisance parameters were regressed out from the time series at each voxel: slow time drifts (basis of discrete cosines with a 0.01 Hz high-pass cut-off), average signals in conservative masks of the white matter, and the lateral ventricles as well as the first principal components of the six rigid-body motion parameters and their squares (Giove et al 2009). Finally, the fMRI volumes were spatially smoothed with a 6 mm isotropic Gaussian blurring kernel and the gray matter (GM) voxels were extracted based on a probabilistic atlas (0.5 was used as the GM probability threshold).

Following this preprocessing, we computed the correlations between the time series of GM voxels which were at least 75 mm apart from each other. We use 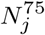 to denote the set of voxels at least 75 mm apart from the voxel *j* and *z*(*r*)_*jj’*_ to denote the Fisher transformed correlation coefficient between the voxels *j* and *j’*. Then, two features 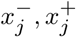 are defined per voxel:

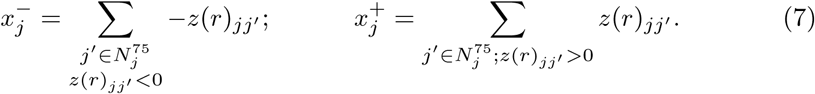

The long-range connection threshold of 75 mm is rather arbitrary, but it has been previously used to define short and long range connections (e.g., by Guo et al (2015); Wang et al (2014)). We separated the positive and negative connections as Guo et al (2015). This preprocessing yielded altogether 81404 variables, corresponding to two times 40702 GM voxels.

## 4. Compared methods

### 4.1 SVM with permutation test

The closest approach to SCBs is training a linear SVM and studying the importance of the weights of the primal variables in the SVM by means of a permutation test (Mouro-Miranda et al 2005; Wang et al 2007). Here, we use two analytic implementations of this approach. The first one involves a null hypothesis test on a statistic over the j-th SVM primal weight (Gaonkar and Davatzikos 2013),*w*_*j.*_ The second one involves a test on its contribution to the SVM margin (Gaonkar et al 2015). As the number of primal variables (voxels) greatly exceeds the number of dual variables (subjects), we can consider that all the training samples would eventually become support vectors. Therefore, we approximate the SVM primal weight vector by the LS-SVM as (Suykens and Vandewalle 1999):

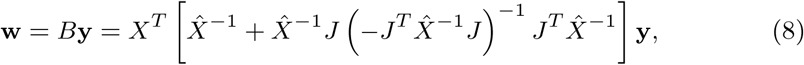

where *X* is the *N* × *P* (number of subjects × number of variables) training data matrix, 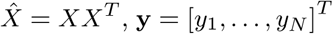 is the associated class label vector, and *J* is a vector of ones. Considering that the permutation test randomly generates different label values with probabilities *P* {*y*_*i*_ = 1} = *p*_1_*, P* {*y*_*i*_ = *-*1} = 1 *- p*_1_ where *p*_1_ is the percentage of patient data, we can define the expected value and variance of the labels during permutations as: 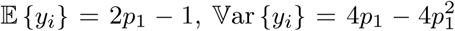. Now, using (8), we define the distribution of the null hypothesis under the first statistic (denoted as SVM+perm) as a Gaussian distribution with mean and variance given by:

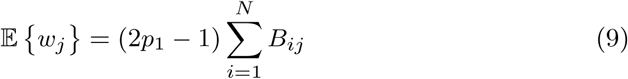

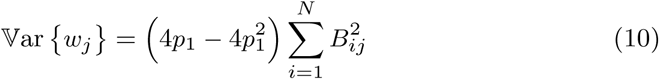

where *B* is as in Eq. (8). The second approach, called SVMmar+perm, defines a statistic over the contribution of the weight of the j-th voxel to the SVM margin as:

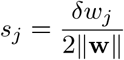

where *δ* is the SVM margin. In this case, its null distribution is approximated by a Gaussian distribution with zero mean and variance

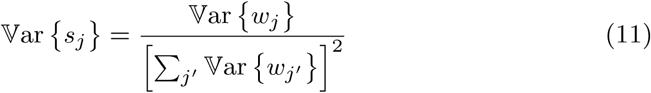

Thus, the test SVM+perm (or SVMmar+perm) will claim that a variable is relevant with a confidence level of *α*, if the probability that a Gaussian distribution, with mean (9) and variance (10) (or zero mean and variance (11)), generates the value *w*_*j*_ (or *s*_*j*_) in the interval 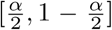. For the experiments we will set the confidence level *α* to 0.05.

### 4.2 T-test and Gaussian Naive Bayes (T-test+NGB)

Although the central part of the discussion is focused on the advantages of SCB over the SVM+perm methodology of previous sub-section, we demonstrate the advantages of SCB over a typical univariate filter-based variable selection/importance. The most widely used massively univariate approach to variable importance is to apply the t-test to each variable separately. Once these tests are applied, the selection of the variables that will be used during the classification can be performed by determining a suitable *α*-threshold on the outcome of the tests, and selecting as important variables those that exceed the corresponding threshold. We use the Gaussian Naive Bayes classifier (John and Langley 1995) as the classifier that consumes the variables selected with the t-test filters. As with the other approaches, we set the *α*-threshold to 0.05, two-sided.

## 5 Results

### 5.1 Synthetic data

Table 1 lists the results achieved on the synthetic data. We evaluated:

- the classification accuracy (ACC) computed using a separate and large test sample;
- the sensitivity (SEN) of the variable selection defined as the ratio between the number of correctly selected important variables and the number of important variables;
- the specificity (SPE) of the variable selection defined as the ratio between the number of correctly identified noisy variables and the number of noisy variables;
- the mean absolute error (MAE) defined as:

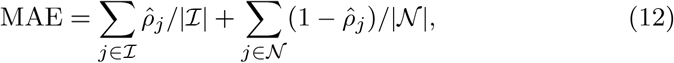

where 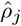 is the estimated p-value for the variable *j* to be important (the lower the p-value the more important the variable), and *ℐ*, *𝒩* are the sets of the important and noise variables, respectively. For the SCB methods, 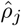 values were computed based on Eq. (6).

**Table 1.** Quantitative results with synthetic data. The values shown are averages and standard deviations over 10 different training sets. ACC is the classification accuracy evaluated using a large test set, SEN is the sensitivity of the variable selection, SPE is the specificity of the variable selection, and MAE is the mean absolute error. See the text for details. Variables are selected using the *α*-threshold of 0.05.

| Method | ACC | SEN | SPE | MAE |
| --- | --- | --- | --- | --- |
| SCB | **0.916** ± 0.004 | **0.369** ± 0.013 | 0.889 ± 0.002 | 0.392 ± 0.004 |
| SCBconf | 0.879 ± 0.008 | 0.208 ± 0.011 | 0.957 ± 0.002 | **0.380** ± 0.006 |
| SVM+perm | 0.797 ± 0.005 | 0.076 ± 0.004 | **0.992** ± 0.001 | 0.411 ± 0.004 |
| SVMmar+perm | 0.891 ± 0.007 | 0.233 ± 0.011 | 0.945 ± 0.001 | 0.411 ± 0.004 |
| t-test+NGB | 0.818 ± 0.010 | 0.259 ± 0.013 | 0.949 ± 0.002 | 0.396 ± 0.004 |

ACC, SEN, SPE measures depend on a categorization of variables into important ones and noise. The categorization, since all the studied methods provide p-values for the variable importance, was determined by a (two-sided) *α*-threshold of 0.05.

Table 1 shows that the accuracies of SCB methods and SVMmar+perm were better than those of the other methods. Indeed, a hypothesis test comparing the accuracies of the methods (t-test, not to be confused with the hypothesis tests about variable importance) over the 10 different training sets indicated a p-value *<* 0.001 in every case. The MAEs achieved by the SCB methods compared very favorably to all baseline approaches (the statistical significance evaluated with t-tests in the 10 data partitions provided a p-value *<* 0.05). Notice that the MAE is independent of the thresholds used to categorize variables as important or not. The specificity (or 1 - SPE) values of the methods are interesting as they can be compared to the nominal *α*-threshold of 0.05; SCB without conformal analysis was too lenient compared to the nominal threshold while the SCBconf well attained the nominal threshold. SVM+perm was overly conservative while the t-test filter and SVMmar+perm attained well the nominal level. The examples in Fig. 1 visualize the same conclusions. Interestingly, as visible in Fig. 1, there was a tendency for all methods to give a high importance to the same variables. This was as expected with a relatively simple simulation.

**Fig. 1.**
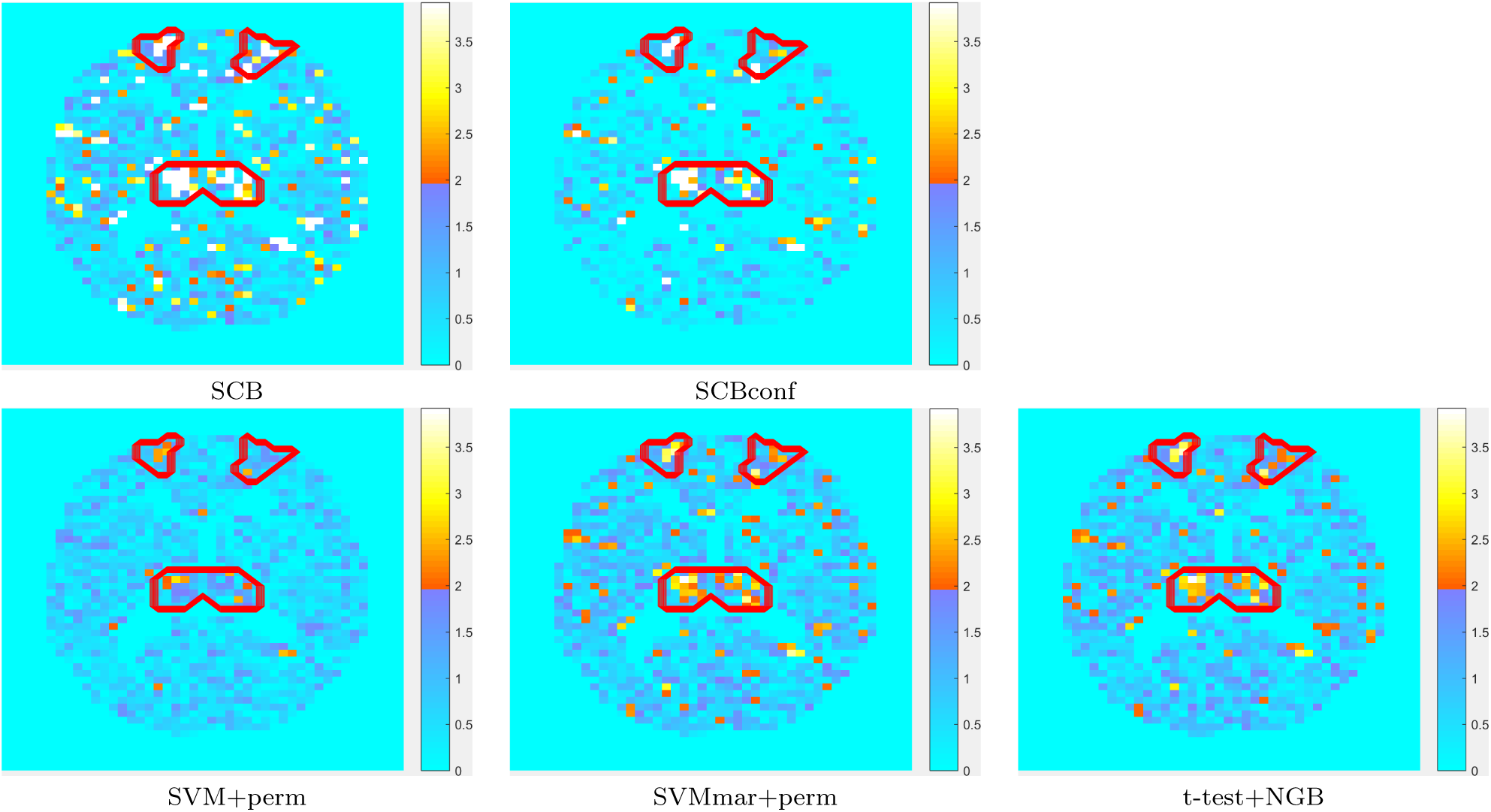
Variable importance *Z*-scores (absolute values) on a plane cutting through Thalami and Superior Frontal Gyri with synthetic data. The voxels in the areas surrounded by red color were important in the ground-truth and the voxels outside those areas were not.

### 5.2 ADNI

With ADNI data, we performed a split-half resampling (2-fold cross-validation) type analysis akin to Tohka et al (2016). This analysis informs us, in addition to the average performance of the methods, about the variability of variable importances due to different subject samples in the same classification problem.

The quantitative results are listed in Table 2. We recorded the test accuracy (ACC) of each algorithm (the fraction of the correctly classified subjects in the test half) averaged across *L* = 100 re-sampling iterations. Moreover, we computed the average absolute difference in ACC between the two split-halves, i.e.,

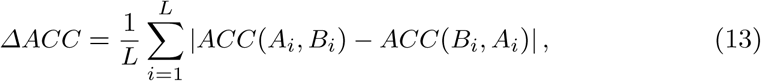

where *ACC*(*A*_*i*_*, B*_*i*_) means accuracy when the training set is *A*_*i*_ and the test set is *B*_*i*_. SCBconf and SCB performed similarly in terms of the classification accuracy and Δ*ACC*. SCB methods were significantly more accurate than t-test+NGB (p-value *<* 0.05) according to a conservative corrected repeated 2-fold CV t-test (Bouckaert and Frank 2004; Nadeau and Bengio 2003) of the classification accuracy. However, this conservative test did not indicate a significant difference between the accuracy of the SCB methods and SVM+perm and SVMmar+perm. Δ*ACC* was markedly smaller with the SCB based methods than with the three other methods.

**Table 2.**
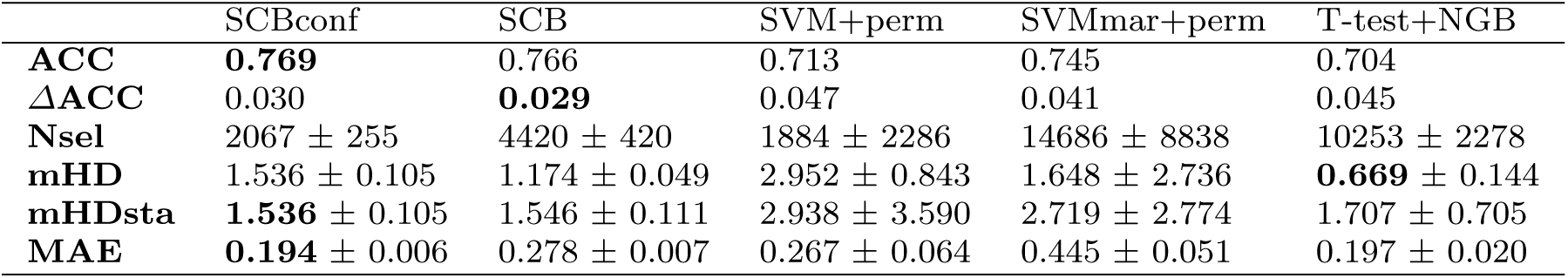
Quantitative results with the ADNI split-half experiment. The values listed are the averaged values over 100 resampling runs followed, where reasonable, by their standard deviations. mHD and mHDsta are computed in voxels. ACC is the classification accuracy, ΔACC is the variability of the ACC (Eq. (13)), Nsel is the number of selected voxels, mHD is the modified Hausdorff distance (Eq. (14)), mHDsta is the modified Hausdorff distance when all methods are forced to select the same number of variables, and MAE is the mean absolute error between the variable importance p-values obtained using independent training sets.

The average number of selected voxels (with *α*-threshold of 0.05) was the smallest with SCBconf and SVM+perm. SCB selected roughly twice as many voxels as SCBconf. The t-test and SVMmar+perm were the most liberal selection methods. When evaluating the standard deviations in the numbers of selected voxels, SCB and SCBconf were the most stable methods. Especially, the number of voxels selected by SVM+perm and SVMmar+perm varied considerably as demonstrated in Fig. 2. We interpret this as a handicap of SVM+perm and SVMmar+perm as the *α*-threshold was always the same and the variability was not expected. The numbers of selected voxels were more variable with T-test+NGB than with SCB-based methods. According to the MAE measure, SCBconf and t-test were the most reproducible (see Table 2).

**Fig. 2.**
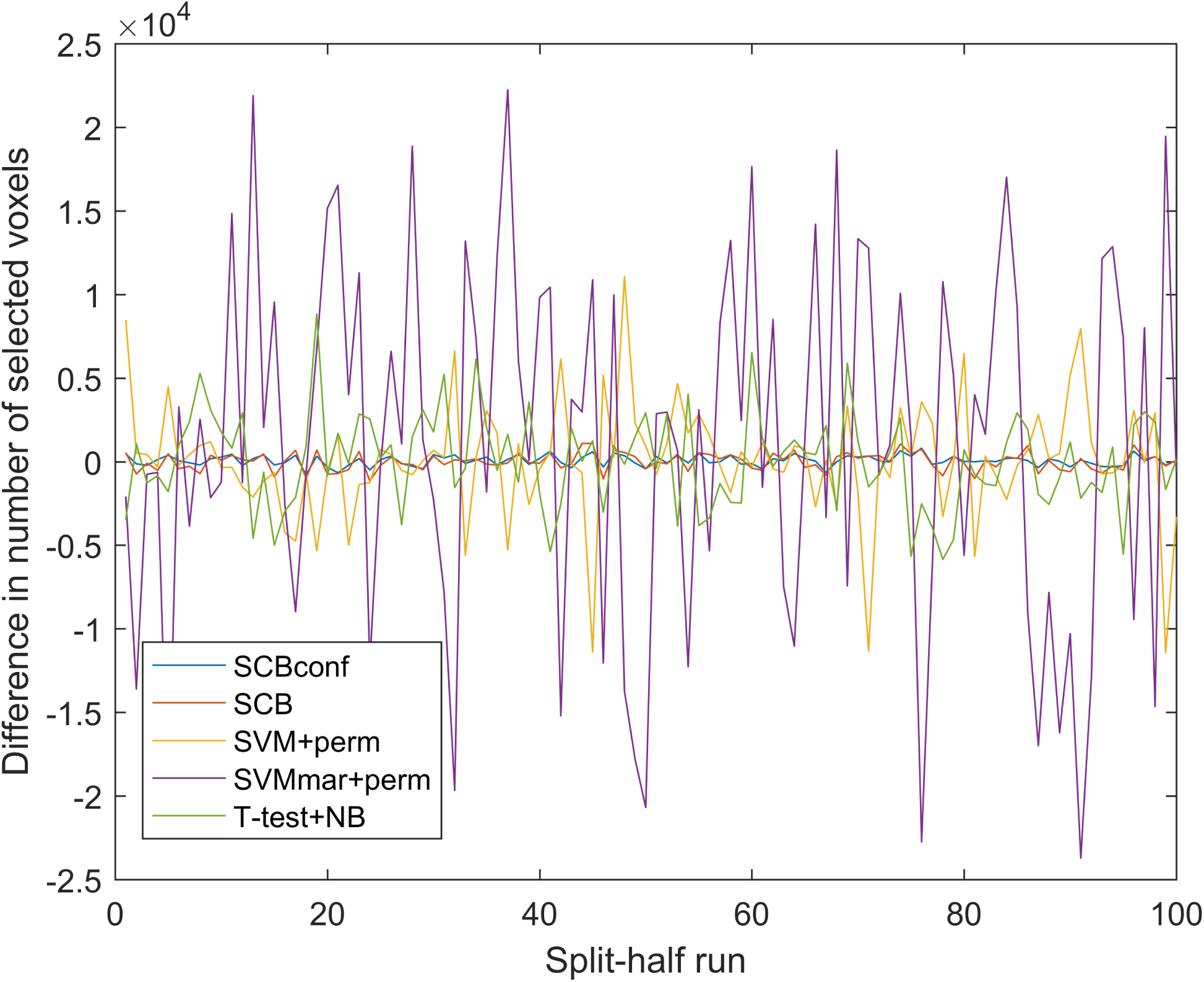
The difference in the numbers of selected voxels between two independent training sets within each split-half resampling run. The SCB methods were more stable with respect to the number of selected voxels than the other methods. Especially, SVM+perm and SVMmar+perm suffered from an excess variability.

We quantified the similarity of two voxel sets selected on the split-halves *A*_*i*_ and *B*_*i*_ using modified Hausdorff distance (mHD) (Dubuisson and Jain 1994). This has the advantage of taking into account spatial locations of the voxels. Let each of the voxels **a** be denoted by its 3-D coordinates (*a*_*x*_*, a*_*y*_*, a*_*z*_). Then, the mHD is defined as

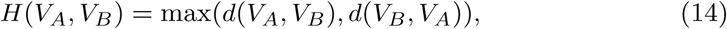

where

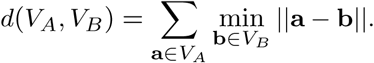

Tohka et al (2016) showed that reproducibility measures of the voxel selection are correlated with the number of selected voxels. To overcome this limitation and make the comparison fair, we here studied standardized sets of voxels by forcing each algorithm to select the same number of voxels as SCBconf in the split half *A*_*i*_. For each algorithm, we then selected the voxels in the *B*_*i*_ according to the *α*-threshold obtained for the split-half *A*_*i*_. The mHD computed using this standardization is denoted by mHDsta in Table 2. As shown in Table 2, the t-test filter was the most reproducible according to the uncorrected mHD. However, this was an artifact of the over-liberality of the test. When standardized with the respect to the number of selected voxels (the row mHDsta), the SCB based methods were most reproducible; however, the difference to the t-test filter was not statistically significant. The SVM+perm and SVMmar+perm were significantly less reproducible than any of the other methods.

Fig. 3 shows examples of visualized variable importance maps. All methods displayed, for example, Hippocampus and Amygdala as important as would be expected. An interesting difference can be observed in middle frontal gyrus, where there was a cluster of highly important voxels according to the SCB methods. However, the t-test filter did not consider these voxels as important. Both SCB methods identified several clusters of important voxels, with SCBconf being more conservative. SVM+perm importance appeared to be more scattered and the t-test was the most liberal selecting many more voxels than the other methods.

**Fig. 3.**
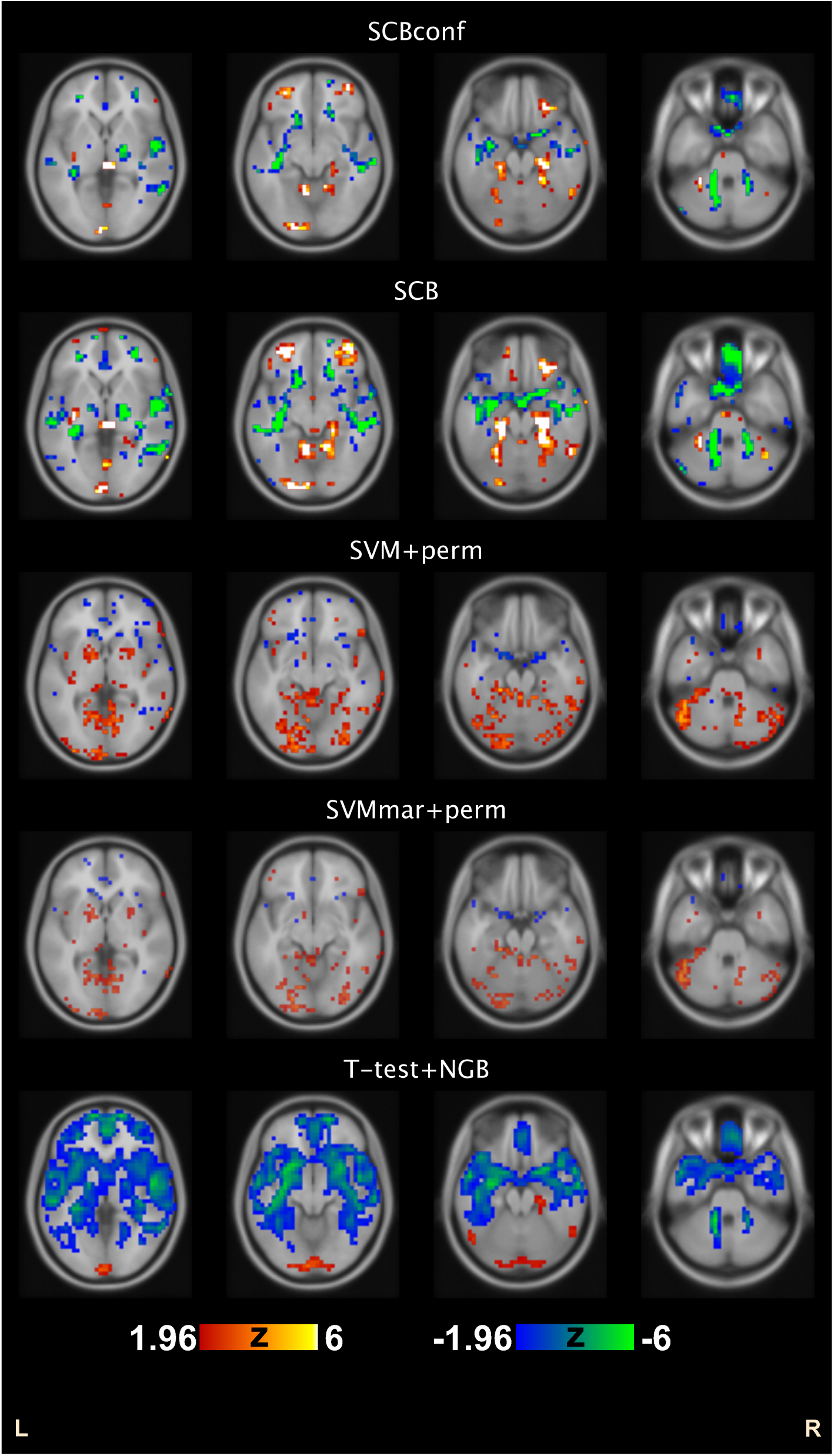
Variable importance Z-scores from a randomly selected example run of the ADNI split-half experiment. The Z-scores are thresholded at |*Z| >* 1.96, corresponding to two-sided alpha threshold of 0.05. Positive *Z* values indicate positive weights. Axial slices at the z-coordinate of the MNI stereotactic space of 0mm, −10mm −20mm, and −30mm are shown.

### 5.3 COBRE

The classification accuracies and numbers of selected voxels with the COBRE data are listed in Table 3. In this experiment, SCBconf was significantly more accurate than the other methods (p-value always *<* 0.01, according to the corrected resampled t-test (Bouckaert and Frank 2004; Nadeau and Bengio 2003)). The other methods performed similarly in terms of the cross-validated classification accuracy. This indicates that the conformal analysis was an essential addition to SCB, probably because the COBRE dataset can be assumed to be more heterogeneous than the ADNI dataset. The heterogeneity of COBRE data probably stems from multiple sources. For example, schizophrenia has often been characterized as a heterogeneous disorder (Seaton et al 2001), the subjects suffering from schizophrenia were receiving various medications at the time of scanning (Kim et al 2016), the age range of the subjects in the dataset was large, and resting state fMRI is more prone to noise due to, for example, subject motion than anatomical T1-weighted MRI. It is particularly in these kinds of applications where we expect the conformal analysis to be most useful. The classification accuracy achieved with SCBconf appeared to outperform recent published analyses of the same data (Chyzhyk et al 2015; Kim et al 2016). However, note that the direct comparison of the classification performance with these works is not fair since it is subject to the differences in variable extraction (different variables were used), data processing (different subjects were excluded) and evaluation (different cross-validation folds were used).

**Table 3.** Average accuracy and number of selected voxels with the 10-fold CV with the COBRE experiment. The values after ± refer to the standard deviations over 10 CV-folds.

|  | SCBconf | SCB | SVM+perm | SVMmar+perm | T-test+NGB |
| --- | --- | --- | --- | --- | --- |
| **ACC** | **$0.952 \pm 0.069$** | $0.695 \pm 0.154$ | $0.731 \pm 0.136$ | $0.692 \pm 0.129$ | $0.709 \pm 0.170$ |
| **Nsel** | $4251 \pm 598$ | $11085 \pm 588$ | $2433 \pm 216$ | $6975 \pm 442$ | $26757 \pm 3397$ |

The SCBconf selected, on average, 4251 variables and was more conservative than the plain SCB, which selected 11085. SVM+perm selected only 2433 variables on average whereas SVMmar+perm selected 6975. The numbers of variables selected by SVM+perm and SVMmar+perm were less variable than in the ADNI experiment where this variation was clearly a problem for these methods. The t-test was overly liberal. Interestingly, the t-test selected many more variables corresponding to the negative correlation strength (on average 24283) than to the positive correlation strength (on average 2474). Instead, SCB and SVM+perm methods selected similar numbers of variables corresponding to the positive and negative correlation strength. This is also visible in Fig. 4, where the median magnitudes of the variable importances are visualized (medians of absolute value of z-scores, see Eq. (6), over 10 CV runs). Concentrating on the SCBconf, widely distributed and partially overlapping areas were found to be important for both negative and positive correlation strength. Particularly, the most important variables (with medians of absolute z-scores exceeding 15 or equivalent p-values smaller than 10^*-*51^) were found in left cerebellum, left inferior temporal gyrus, left and right thalamus, left inferior parietal gyrus, right inferior frontal gyrus, left medial frontal gyrus, and left middle frontal gyrus for negative correlation strength. For positive correlation strength, median absolute z-scores exceeding 15 were found in left and right cerebellum, left inferior frontal gyrus, left caudate, right lingual gyrus, right middle temporal gyrus and left medial frontal gyrus. We note that a high z value of 15 was selected as threshold in this discussion to concentrate only to the most important variables. We have made the complete maps of variable importance available at NeuroVault serviceGorgolewski et al (2015) at http://neurovault.org/collections/MOYIOPDI/.

**Fig. 4.**
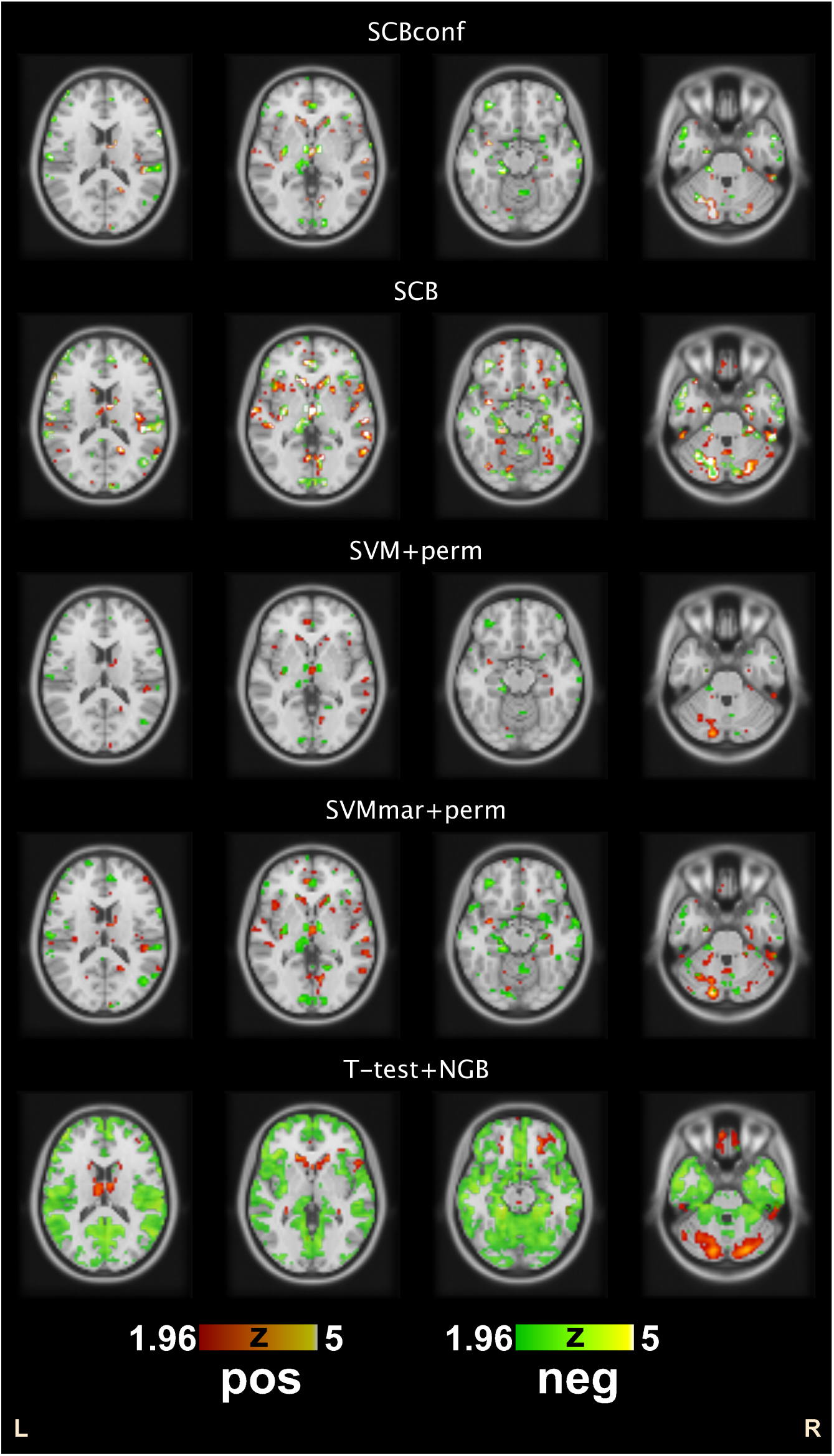
Median magnitudes of variable importance Z scores among 10 CV runs with COBRE data. The Z-scores are thresholded at |*Z| >* 1.96. Note that if a variable lights up then it was selected during at least half of the CV runs. ‘Pos’ and ‘Neg’ quantifiers refer to the strength of the positive and negative connectedness that were separated in the analysis. We do not visualize whether the classifier weights are negative or positive to avoid clutter. Axial slices at the z-coordinate of the MNI stereotactic space of 15mm, 0mm −15mm, and −30mm are shown. Complete maps are available in the NeuroVault service http://neurovault.org/collections/MOYIOPDI/.

With the COBRE data, we studied the effect of multiple comparisons correction to the classification accuracy and to the number of selected variables. For multiple comparisons correction, we used variable-wise false discovery rate (FDR) correction with Benjamini-Hochberg procedure (assuming independence) (Benjamini and Hochberg 1995). The classification accuracies and the numbers of selected variables, with and without FDR correction, are shown in box-plots of Figure 5. SVM+perm and SVMmar+perm were excluded from this experiment as the multiple comparisons problem is different with them (Gaonkar and Davatzikos 2013) and it was found to produce an empty set of variables in some cases. As is shown in Figure 5, including multiple comparisons correction had no influence to the classification performance with any of the methods.

**Fig. 5.**
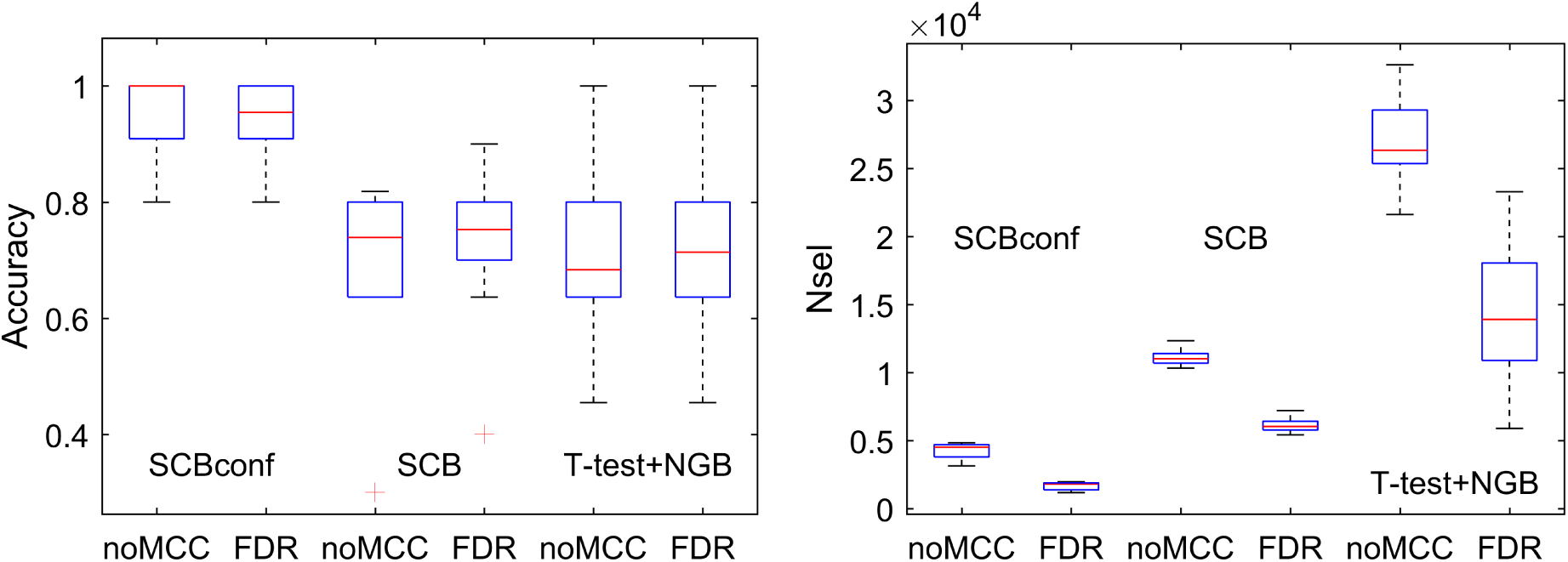
The classification accuracy and the number of selected variables across 10 CV folds with COBRE data with and without FDR based multiple comparisons correction. Whether FDR correction is included or not made no difference to the classification performance of the methods.

### 5.4 Computation time

The experiments were run in a computer with Intel Xeon 2.40Ghz processor with 20 cores and 128 Gb of RAM. The training of several SVMs that took place in the bagging stages of both SCB and SCBconf was parallelized across all the cores of the computer. All the other computations (the weight aggregation that leads to the final measure of variable importance, the hypothesis testing and the evaluation of the final SVM) were done using a single core. The baseline methods, SVM+perm, SVMmar+perm, and t-test+ NGB, were run using a single core.

The baseline methods SVM+perm, SVMmar+perm, and t-test+ NGB required between 1 and 5 seconds depending on the size of the dataset (number of samples and dimensionality) and on the number of selected important variables, as this last quantity determines the training time of the final classifier. The computation time of the SCB was in the range 5 to 6 minutes due to the bagging. The computation time was up to 2 hours in the case of the SCBconf, as each conformal analysis iteration involves a complete bagging and we carried out *R* = 20 of these iterations. Since bagging can be run in parallel, these times could be substantially reduced by further parallelization.

## 6. Discussion

We have introduced and evaluated new variable importance measures, termed SCB and SCBconf, based on sign consistency of classifier ensembles. The measures are specially designed for very high dimensional scenarios with far more variables than samples such as in neuroimaging, where many widely used multivariate variable importance measures fail. The SCB variable importance measures extend and generalize ideas for the voxel selection we have introduced earlier (Parrado-Hernández et al 2014). We have derived a novel parametric hypothesis test that can be used to assign a p-value to the importance of the variable for a classification. We have shown that the variable selection using SCB and SCBconf importance measures leads to a more accurate classification than the variable selections based on a standard massively univariate hypothesis testing, or on two statistical tests based on a parametric SVM permutation test (Gaonkar and Davatzikos 2013; Gaonkar et al 2015). These three methods were compared to the SCB methods because 1) they are applicable to high-dimensional data and 2) they come with a parametric hypothesis test to assign p-values to variable importance. We have also demonstrated that the proposed SCB and SCBconf measures were robust and that they can lead to better classification accuracies than the state of art in schizophrenia classification based on resting state fMRI.

The basic idea behind the SCB methods is to train several thousand linear SVMs, each with a different subsample of data, and then study the sign-consistency of weights assigned to each variable. These weights having the same sign is a strong indication of the stability of the interpretation of the variable with respect to random subsampling of the data and, thus, a strong indication of the importance of the variable. Therefore, we can quantify the importance of the variable by studying the frequency of sign of the classifier weights assigned to it. While the ideas of random subsampling and random relabeling are widely used for variable importance and selection, for example, in the out-of-bag variable importances of RFs (Breiman 2001), the idea of sign consistency is much less exploited and novel in brain imaging. The reason to study signs of the weights of linear SVMs rather magnitudes of those weights is that the signs of the weights are much less sensitive to the redundancy of the data (multicolinearity). In addition, the signs of the weights have a simple interpretation in terms of the classification task while the weight magnitude is much more difficult to interpret in high dimensions. A positive weight means that the associated variable pushes the classifier towards the positive class while a negative weight means that the associated variable pushes the classifier towards negative class. The conformal variant, SCBconf, refines SCB variable importance by utilizing test data by assigning random labels to it. This is essentially relabeling in the transductive setting and it is especially useful in situations where the the data is heterogeneous as we demonstrated using the COBRE resting-state fMRI sample. The reason why the refinement is needed is that most variables in a brain scan contribute to separating that one brain from the other brains in the training set. Expressed in terms of linear classification, the number of linearly independent components among the training input variables exceeds the number of training instances so that any set of binary labels leads to a linearly separable problem, and randomizing a part of the labeling helps in separating truly important variables for the characterization of the disease from the variables that just separate one brain from the rest.

We note that the SCB variable importances are probabilistic, i.e., if the signs of the weights of classifiers from the bagging process are consistent with a high probability, then the associated variable is deemed as important for the classification. The nature of the quantifier ‘with a high probability’ is made exact by the associated p-value resulting from the hypothesis test against the null hypothesis of that the sign is inconsistent. We further note that even for the most important variables, two subsets of training data leading to opposite signs for weight of that variable probably exist. However, for the important variable, the signs are the same with high probability.

We have used uncorrected p-values to threshold the variable importance scores. There are two reasons for this. First, the variable importance scores might be interesting also for variables that do not pass stringent multiple comparisons corrected threshold. Second, retaining also variables that are borderline important could improve the generalization performance of the classifier. With the COBRE fMRI dataset, we have shown that ultimately this is a matter of preference and whether using corrected or uncorrected thresholds makes no difference to the generalization performance of the classifier. We also experimented this with synthetic data and observed a slight drop in the classification performance when using the FDR corrected thresholds. As Gaonkar and Davatzikos (2013) noted the classifier weights of an SVM are not independent and thus FDR based multiple comparisons correction probably over-corrects. In a data-rich situation, cross-validation based estimate of the generalization error might be used to select the optimal *α*-threshold, however, one should keep in mind that cross-validation based error estimates have large variances (Dougherty et al 2011) and this might offset the potential gains of not setting the importance threshold a-priori (Tohka et al 2016; Huttunen and Tohka 2015; Varoquaux et al 2017).

The SCB method has two parameters: the number of resampling iterations *S* and the subsampling rate *γ*. In our target applications, where the number of variables is larger than the number of samples, the parameter *C* for the SVMs can always be selected to be large enough (here *C* = 100) to ensure full separation. For the parameter *S*, the larger value is always better and we have found that *S* = 10.000 has been sufficient. We have selected the subsampling rate to be 0.5 and previously we have found that the method is not sensitive to this parameter; in fact, these parameter settings agree with those previously used by Parrado-Hernández et al (2014). SCBconf has one extra parameter *R* (the number of random labelings of the test samples). We have here selected *R* = 20 and we do not expect gains by increasing this value.

It is important to compare our approach to RFs (Breiman 2001), which also use bagging to derive variable importances from several classifiers. An important difference is the choice of the base classifier used to construct an ensemble. In our case, this is a linear classifier, where each variable receives a different weight in each classifier of the ensemble. These different weights received by a same variable across all the classifiers in the ensemble can be combined in a score that decides the importance of the variable. RFs, instead, use decision trees as the base classifiers. As we have argued in the introduction, the use of linear classifiers is advantageous when the number of variables is in the order of tens of thousands as the number of trees in a forest would have to be extraordinarily large to decide importance for each variable. We also remark that when the number of samples is smaller than the number of variables, the classification tasks are necessarily linearly separable and thus solvable by linear classifiers (Duda et al 2012). Techniques such as oblique RFs (oRFs) (Menze et al 2011), offer an interesting half way between RFs based on univariate splits and linear classifiers as in each node the decision is based on a linear classifier with a randomly selected subset of variables (Menze et al (2011) suggest to use subsets of the size 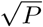). In the cases where subsets of variables used for training a classifier is larger than the number of training subjects, each tree in an oRF reduces to a single linear classifier because the classification problem in the first node is already linearly separable and thus the subsequent nodes would face data subsets where all the instances belong to the same class. Variable importance in oRFs depend on the choice of an additional tuning parameter to decide if a variable is important in an individual split or not.

Placing a p-value to variable importances in RFs has turned out to be a difficult problem. The early proposal by Breiman and Cutler (2007) suffers from substantial problems identified by Strobl and Zeileis (2008). Later proposals (Hapfelmeier and Ulm 2013; Altmann et al 2010) rely on permutation approaches requiring training hundreds of RFs even when the number of variables is in the order of tens (Hapfelmeier and Ulm 2013). Consequently, variable selection approaches using RFs rely on different heuristics or cross-validation (Díaz-Uriarte and De Andres 2006; Genuer et al 2010).

### Information Sharing Statement

Only open datasets were used in the paper. The Alzheimer’s Disease Neuroimaging Initiative (ADNI) is available at http://www.adni-info.org. The pre-processed form of the COBRE dataset is available at https://figshare.com/articles/COBRE_preprocessed_with_NIAK_0_12_4/1160600. The original COBRE dataset can be found in International Neuroimaging Data-sharing Initiative (INDI)^8^. The software implementation of all the methods has been developed in Python and a Python notebook containing the implementations is https://github.com/vgverdejo/ResearchActivities/blob/master/Neuroimage/Sign-consistency.ipynb. Statistical maps are available at http://neurovault.org/collections/MOYIOPDI/.

## Supporting information

## Acknowledgments

Data collection and sharing for this project was funded by the Alzheimer’s Disease Neuroimaging Initiative (ADNI) (National Institutes of Health Grant U01 AG024904) and DOD ADNI (Department of Defense award number W81XWH-12-2-0012). ADNI is funded by the National Institute on Aging, the National Institute of Biomedical Imaging and Bioengineering, and through generous contributions from the following: AbbVie, Alzheimer’s Association; Alzheimer’s Drug Discovery Foundation; Araclon Biotech; BioClinica, Inc.; Biogen; Bristol-Myers Squibb Company; CereSpir, Inc.; Eisai Inc.; Elan Pharmaceuticals, Inc.; Eli Lilly and Company; EuroImmun; F. Hoffmann-La Roche Ltd and its affiliated company Genentech, Inc.; Fujirebio; GE Healthcare; IXICO Ltd.; Janssen Alzheimer Immunotherapy Research & Development, LLC.; Johnson & Johnson Pharmaceutical Research & Development LLC.; Lumosity; Lundbeck; Merck & Co., Inc.; Meso Scale Diagnostics, LLC.; NeuroRx Research; Neurotrack Technologies; Novartis Pharmaceuticals Corporation; Pfizer Inc.; Piramal Imaging; Servier; Takeda Pharmaceutical Company; and Transition Therapeutics.

J. Tohka’s work was supported by the Academy of Finland (grant no. 316258) and V. Gómez-Verdejo’s work has been partly funded by the Spanish MINECO projects TEC2014-52289R and TEC2016-81900-REDT/AEI.

## Footnotes

1 In most scenarios relevant to this work, a single variable corresponds to a single voxel, but this does not have to be the case. We opt to use the general term variable here.

2 We refer to those RFs in which each tree split is defined in terms of a single variable.

3 Transductive algorithms can be contrasted with inductive ones, for example, the k-nearest neighbor algorithms are transductive algorithms (Gammerman et al 1998). Because the label information of the test data is not used in the learning process, transduction does not lead to upward biased classifier performance estimates.

4 See a Python notebook for examples https://github.com/vgverdejo/ResearchActivities/blob/master/Neuroimage/Sign-consistency.ipynb

5 Available at https://figshare.com/articles/COBRE_preprocessed_with_NIAK_0_12_4/1160600

6 http://fcon_1000.projects.nitrc.org/indi/retro/cobre.html

7 https://github.com/SIMEXP/niak

8 http://fcon_1000.projects.nitrc.org/indi/retro/cobre.html

## References

Altmann A, Toloşi L, Sander O, Lengauer T (2010) Permutation importance: a corrected feature importance measure. Bioinformatics 26(10):1340–1347

Archer KJ, Kimes RV (2008) Empirical characterization of random forest variable importance measures. Computational Statistics & Data Analysis 52(4):2249–2260

Bellec P, Benhajali Y, Carbonell F, Dansereau C, Albouy G, Pelland M, Craddock C, Collignon O, Doyon J, Stip E, Orban P (2015) Impact of the resolution of brain parcels on connectome-wide association studies in fmri. NeuroImage 123:212 – 228, DOI http://dx.doi.org/10.1016/j.neuroimage.2015.07.071, URL http://www.sciencedirect.com/science/article/pii/S1053811915006916

Benjamini Y, Hochberg Y (1995) Controlling the false discovery rate: a practical and powerful approach to multiple testing. Journal of the royal statistical society Series B (Methodological) pp 289–300

Bi J, Bennett K, Embrechts M, Breneman C, Song M (2003) Dimensionality reduction via sparse support vector machines. JMLR 3:1229–1243

Bouckaert RR, Frank E (2004) Evaluating the replicability of significance tests for comparing learning algorithms. In: Advances in knowledge discovery and data mining, Springer, pp 3–12

Breiman L (2001) Random forests. Machine Learning 45(1):5–32

Breiman L, Cutler A (2007) Random forests-classification description. Department of Statistics, Berkeley

Caragea D, Cook D, Honavar VG (2001) Gaining insights into support vector machine pattern classifiers using projection-based tour methods. In: Proceedings of the seventh ACM SIGKDD international conference on Knowledge discovery and data mining, ACM, pp 251–256

Chang CC, Lin CJ (2011) Libsvm: a library for support vector machines. ACM Transactions on Intelligent Systems and Technology (TIST) 2(3):27

Chu C, Hsu AL, Chou KH, Bandettini P, Lin C, Initiative ADN, et al (2012) Does feature selection improve classification accuracy? impact of sample size and feature selection on classification using anatomical magnetic resonance images. Neuroimage 60(1):59–70

Chyzhyk D, Savio A, Graña M (2015) Computer aided diagnosis of schizophrenia on resting state fmri data by ensembles of elm. Neural Networks 68:23–33

Cohen JR, Asarnow RF, Sabb FW, Bilder RM, Bookheimer SY, Knowlton BJ, Poldrack RA (2010) Decoding developmental differences and individual variability in response inhibition through predictive analyses across individuals. The developing human brain p 136

Collins DL, Evans AC (1997) Animal: validation and applications of nonlinear registration-based segmentation. International journal of pattern recognition and artificial intelligence 11(08):1271–1294

Díaz-Uriarte R, De Andres SA (2006) Gene selection and classification of microarray data using random forest. BMC bioinformatics 7(1):3

Dougherty E, Zollanvari A, Braga-Neto U (2011) The illusion of distribution-free small-sample classification in genomics. Current genomics 12(5):333–341

Dubuisson MP, Jain AK (1994) A modified hausdorff distance for object matching. In: Pattern Recognition, 1994. Vol. 1-Conference A: Computer Vision & Image Processing., Proceedings of the 12th IAPR International Conference on, IEEE, vol 1, pp 566–568

Duda RO, Hart PE, Stork DG (2012) Pattern classification. John Wiley & Sons

Dukart J, Schroeter ML, Mueller K, Initiative ADN, et al (2011) Age correction in dementia–matching to a healthy brain. PloS one 6(7):e22, 193

Fonov V, Evans AC, Botteron K, Almli CR, McKinstry RC, Collins DL, Group BDC, et al (2011) Unbiased average age-appropriate atlases for pediatric studies. NeuroImage 54(1):313–327

Friedman J, Hastie T, Tibshirani R (2008) The elements of statistical learning 2nd Ed., vol 1. Springer series in statistics Springer, Berlin

Gammerman A, Vovk V, Vapnik V (1998) Learning by transduction. In: AISTATS98, Morgan Kaufmann Publishers Inc., pp 148–155

Gaonkar B, Davatzikos C (2013) Analytic estimation of statistical significance maps for support vector machine based multivariate image analysis and classification. NeuroImage 78:270–283

Gaonkar B, Shinohara RT, Davatzikos C (2015) Interpreting support vector machine models for multivariate group wise analysis in neuroimaging. Medical Image Analysis 24(1):190–204, DOI https://doi.org/10.1016/j.media.2015.06.008, URL http://www.sciencedirect.com/science/article/pii/S136184151500095X

Gaser C, Franke K, Klöppel S, Koutsouleris N, Sauer H, Initiative ADN, et al (2013) BrainAGE in mild cognitive impaired patients: predicting the conversion to alzheimer’s disease. PloS ONE 8(6):e67,346

Genuer R, Poggi JM, Tuleau-Malot C (2010) Variable selection using random forests. Pattern Recognition Letters 31(14):2225–2236

Giove F, Gili T, Iacovella V, Macaluso E, Maraviglia B (2009) Images-based suppression of unwanted global signals in resting-state functional connectivity studies. Magnetic resonance imaging 27(8):1058–1064

Gomez-Verdejo V, Parrado-Hernandez E, Tohka J (2016) Voxel importance in classifier ensembles based on sign consistency patterns: application to smri. In: Pattern Recognition in Neuroimaging (PRNI), 2016 International Workshop on, IEEE, pp 1–4

Gorgolewski KJ, Varoquaux G, Rivera G, Schwarz Y, Ghosh SS, Maumet C, Sochat VV, Nichols TE, Poldrack RA, Poline JB, et al (2015) Neurovault. org: a web-based repository for collecting and sharing unthresholded statistical maps of the human brain. Frontiers in neuroinformatics 9:8

Greenstein D, Malley JD, Weisinger B, Clasen L, Gogtay N (2012) Using multivariate machine learning methods and structural mri to classify childhood onset schizophrenia and healthy controls. Front Psychiatry 3:53

Grosenick L, Klingenberg B, Katovich, K B Knutson, Taylor JE (2013) Interpretable wholebrain prediction analysis with graphnet. NeuroImage 72:304–321

Guo W, Liu F, Xiao C, Liu J, Yu M, Zhang Z, Zhang J, Zhao J (2015) Increased short-range and long-range functional connectivity in first-episode, medication-naive schizophrenia at rest. Schizophrenia Research 166(1–3):144–150, DOI http://dx.doi.org/10.1016/j.schres.2015.04.034, URL http://www.sciencedirect.com/science/article/pii/S0920996415002297

Guyon I, Weston J, Barnhill S, Vapnik V (2002) Gene selection for cancer classification using support vector machines. Machine Learning 46(1):389–422, DOI 10.1023/A:1012487302797, URL http://dx.doi.org/10.1023/A:1012487302797

Hapfelmeier A, Ulm K (2013) A new variable selection approach using random forests. Computational Statistics & Data Analysis 60:50–69

Haufe S, Meinecke F, Görgen K, Dähne S, Haynes JD, Blankertz B, Bießmann F (2014) On the interpretation of weight vectors of linear models in multivariate neuroimaging. Neuroimage 87:96–110

Huttunen H, Tohka J (2015) Model selection for linear classifiers using bayesian error estimation. Pattern Recognition 48(11):3739–3748

John GH, Langley P (1995) Estimating continuous distributions in bayesian classifiers. In: Proceedings of the Eleventh conference on Uncertainty in artificial intelligence, Morgan Kaufmann Publishers Inc., pp 338–345

Kerr WT, Douglas PK, Anderson A, Cohen MS (2014) The utility of data-driven feature selection: Re: Chu et al. 2012. NeuroImage 84:1107–1110

Khundrakpam BS, Tohka J, Evans AC (2015) Prediction of brain maturity based on cortical thickness at different spatial resolutions. NeuroImage 111:350–359

Kim J, Calhoun VD, Shim E, Lee JH (2016) Deep neural network with weight sparsity control and pre-training extracts hierarchical features and enhances classification performance: Evidence from whole-brain resting-state functional connectivity patterns of schizophrenia. NeuroImage 124:127–146

Langs G, Menze BH, Lashkari D, Golland P (2011) Detecting stable distributed patterns of brain activation using gini contrast. NeuroImage 56(2):497–507

Menze BH, Kelm BM, Splitthoff DN, Koethe U, Hamprecht FA (2011) On oblique random forests. In: Joint European Conference on Machine Learning and Knowledge Discovery in Databases, Springer, pp 453–469

Michel V, Gramfort A, Varoquaux G, Eger E, Thirion B (2011) Total variation regularization for fmri-based prediction of behavior. IEEE transactions on medical imaging 30(7):1328–1340

Moradi E, Pepe A, Gaser C, Huttunen H, Tohka J (2015) Machine learning framework for early mri-based alzheimer’s conversion prediction in mci subjects. Neuroimage 104:398–412

Mouro-Miranda J, Bokde A, Born C, Hampel H, Stetter M (2005) Classifying brain states and determining the discriminating activation patterns: Support vector machine on functional MRI data. NeuroImage 28:980–995

Mwangi B, Tian TS, Soares JC (2014) A review of feature reduction techniques in neuroimaging. Neuroinformatics 12(2):229–244

Nadeau C, Bengio Y (2003) Inference for the generalization error. Machine Learning 52(3):239–281

Parrado-Hernández E, Gómez-Verdejo V, Martínez-Ramón M, Shawe-Taylor J, Alonso P, Pujol J, Menchón JM, Cardoner N, Soriano-Mas C (2014) Discovering brain regions relevant to obsessive–compulsive disorder identification through bagging and transduction. Medical image analysis 18(3):435–448

Pedregosa F, Varoquaux G, Gramfort A, Michel V, Thirion B, Grisel O, Blondel M, Prettenhofer P, Weiss R, Dubourg V, et al (2011) Scikitlearn: Machine learning in python. Journal of Machine Learning Research 12(Oct):2825–2830

Power JD, Barnes KA, Snyder AZ, Schlaggar BL, Petersen SE (2012) Spurious but systematic correlations in functional connectivity mri networks arise from subject motion. Neuroimage 59(3):2142–2154

Seaton BE, Goldstein G, Allen DN (2001) Sources of heterogeneity in schizophrenia: the role of neuropsychological functioning. Neuropsychology review 11(1):45–67

Strobl C, Zeileis A (2008) Danger: High power! – exploring the statistical properties of a test for random forest variable importance. In: Brito P (ed) Proceedings of the 18th International Conference on Computational Statistics, Porto, Portugal (CD-ROM), Springer, pp 59–66

Strobl C, Boulesteix AL, Kneib T, Augustin T, Zeileis A (2008) Conditional variable importance for random forests. BMC bioinformatics 9(1):307

Suykens J, Vandewalle J (1999) Least squares support vector machine classifiers. Neural Processing Letters 9(3):293–300, DOI 10.1023/A:1018628609742, URL https://doi.org/10.1023/A:1018628609742

Tohka J, Moradi E, Huttunen H (2016) Comparison of feature selection techniques in machine learning for anatomical brain mri in dementia. Neuroinformatics 14:279 – 296

Tzourio-Mazoyer N, Landeau B, Papathanassiou D, Crivello F, Etard O, Delcroix N, Mazoyer B, Joliot M (2002) Automated anatomical labeling of activations in spm using a macroscopic anatomical parcellation of the mni mri single-subject brain. Neuroimage 15(1):273–289

Varoquaux G, Raamana PR, Engemann DA, Hoyos-Idrobo A, Schwartz Y, Thirion B (2017) Assessing and tuning brain decoders: cross-validation, caveats, and guidelines. NeuroImage 145:166–179

Wang X, Xia M, Lai Y, Dai Z, Cao Q, Cheng Z, Han X, Yang L, Yuan Y, Zhang Y, Li K, Ma H, Shi C, Hong N, Szeszko P, Yu X, He Y (2014) Disrupted resting-state functional connectivity in minimally treated chronic schizophrenia. Schizophrenia Research 156(2–3):150–156, DOI http://dx.doi.org/10.1016/j.schres.2014.03.033, URL http://www.sciencedirect.com/science/article/pii/S0920996414001728

Wang Z, Childress A, Wang J, Detre J (2007) Support vector machine learning-based fMRI data group analysis. NeuroImage 36:1139–1151

Zou H, Hastie T (2005) Regularization and variable selection via the elastic net. Journal of the Royal Statistical Society: Series B (Statistical Methodology) 67(2):301–320

