## Supplementary material for "Sign-consistency based variable importance for machine learning in brain imaging"

### 1 An example of sign-consistency computations

We demonstrate the computations involved in our method, Sign-consistency bagging (SCB), with a toy example. This example only depicts the use of the basic method, SCB, and it does not depict the use of SCBconf.

Figure 1 shows a dataset proposed by an anonymous reviewer. The problem is linearly separable, in fact, a linear SVM with $C \geq 10^7$ separates the training data with zero errors. Both variables are important for the classification in this example.

To use the proposed method, we need to fix the size of the subset of the training data that will result of the sampling at each iteration of the bagging. We select 3 samples since this number in 2 dimensional datasets guarantees linear separability in all the training subsets (even in the case that the whole dataset would not be linearly separable). Moreover, this number potentially also provides a richer distribution of separating hyperplanes among the members of the ensemble as the distribution of the training subset in each iteration of the ensemble can be very different from the data distribution of the whole dataset.

The use of 3 training examples (from a total of 14 samples) per member of the ensemble brings out a total number of 288 different non-trivial classification datasets. We mean by non-trivial that there is at least one sample from each of the two classes.

We have trained those 288 classifiers with a linear SVM with $C = 10^7$ and obtained the separating planes depicted in Figure 2. White circles in the plots are samples not selected as part of the training set for each particular bagging iteration. One can notice that in a majority of cases both weights are positive. The exact number of times this occurs is the following:

- Weight in coordinate x:
  - Positive 90.97%
  - Zero 1.39%
  - Negative 7.64%

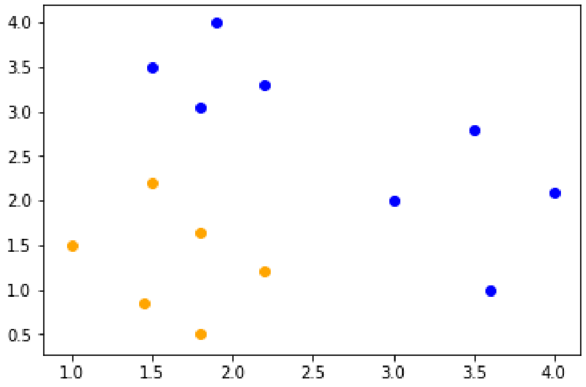


Figure 1: Data set proposed by an anynomous reviewer. The data are split in two classes: blue with 8 samples and orange with 6 samples. It is easy to visually check that the problem is linearly separable.

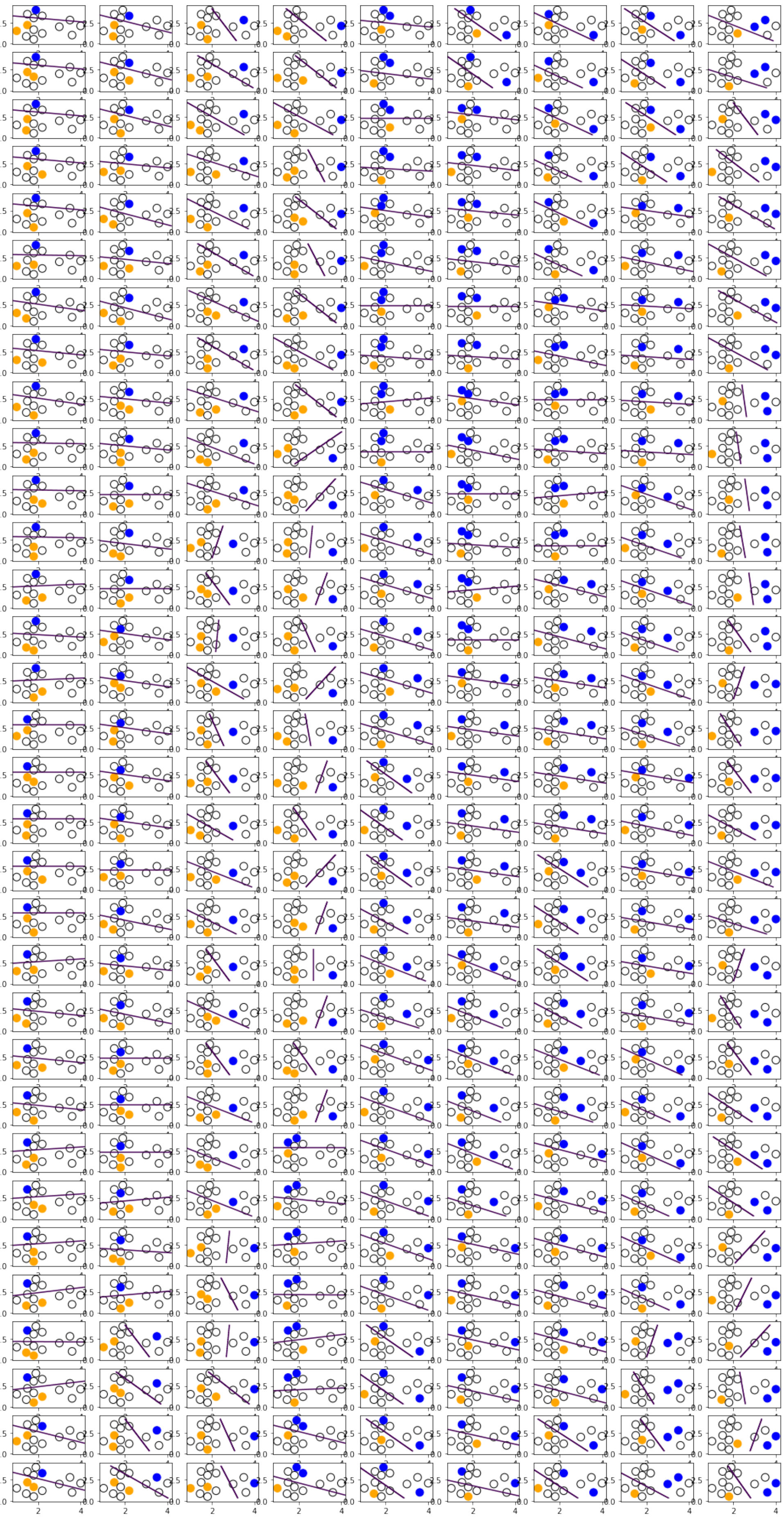

Figure 2: All possible separating planes that could take part in the ensemble that would solve the dataset in Figure 1 of this supplement.

- Weight in coordinate y:
  - Positive 92.36%
  - Zero 0.00%
  - Negative 7.64%

These results give, in the notation of the paper, $\hat{p}_x = 0.910$, $\hat{p}_y = 0.924$, $I_x = 0.820$, $I_x = 0.847$. Further, $\gamma = 3/14$ and $S = 288$, which lead to $\hat{z}_x = 2.74$ and $\hat{z}_y = 3.05$, which indicates that both variables are important according to the hypothesis test, with $p$-values smaller than 0.01.

Although in a number of members of the ensemble the weights can have negative values, there is a high probability that the ensemble captures the actual consistency patterns in the sign of the weights. Notice that in this problem the number of samples is 14 and we have construct an ensemble using only 3 samples to train each member of the ensemble (each training subset size is less that 1/3 of the size of the complete training set size), while in the results presented in the paper the size of the training subsets is 1/2 of the size of the whole set. This results in training subsets being more representative of the actual data distribution but diverse enough to boost the capabilities of the presented method.
